# Cigarette smoking promotes the spread of antimicrobial resistance in the human lung and the environment

**DOI:** 10.1101/2023.08.14.553211

**Authors:** Peiju Fang, Diala Konyali, Emily Fischer, Robin Pascal Mayer, Jin Huang, Alan Xavier Elena, Gerit Hartmut Orzechowski, Andrew Tony-Odigie, David Kneis, Alexander Dalpke, Peter Krebs, Bing Li, Thomas U. Berendonk, Uli Klümper

**Affiliations:** Technische Universität Dresden, Institute of Hydrobiology, Dresden, Germany; Technische Universität Dresden, Institute of Urban and Industrial Water Management, Dresden, Germany; Tsinghua University, Tsinghua Shenzhen International Graduate School, Institute of Environment and Ecology, Shenzhen, China; Technische Universität Dresden, Institute of Medical Microbiology and Virology, University Hospital Carl Gustav Carus, Dresden, Germany; University Hospital Heidelberg, Department of Infectious Diseases, Medical Microbiology and Hygiene, Heidelberg, Germany

**Keywords:** Cigarette smoke condensate, cigarette ash, cigarette filter, plasmid transfer, antibiotic resistance, cigarette smoking

## Abstract

While immediate health risks of cigarette smoking are well-established, indirect health impacts of cigarette-derived pollutants through proliferation of antimicrobial resistance (AMR) among bacteria remain understudied. Here, exposure to cigarette smoke condensate at relevant concentrations resulted in >2-fold elevated transfer rates of a multi-drug-resistance encoding plasmid between *Pseudomonas* strains in artificial lung sputum medium. This effect was connected to elevated reactive oxygen species production as part of the bacterial stress response when exposed to cigarette-derived toxicants. Similar results were obtained under exposure to cigarette ash leachate in environmental medium. Further, used cigarette filters enriched in toxic residues were submerged in a wastewater stream, and colonized by altered microbial communities compared to unused filters. These communities were significantly enriched in pathogens and AMR. Hence, filters could facilitate hitchhiking of high-risk bacteria to novel environments. We demonstrate that cigarette-derived compounds can promote the spread of AMR within the human lung and natural environments.

## Introduction

Antimicrobial resistance (AMR) as well as smoking of tobacco products have been identified as two of the major threats to global human health, with both being associated with millions of deaths every year^1,2^. However, the immediate interactions between these two have rarely been investigated.

Around one in five citizens worldwide smokes cigarettes, with approximately 5.5 trillion cigarettes consumed in 2016, a number that is expected to reach 9 trillion by 2025^3,4^. Tobacco comprises more than 7,000 chemical compounds and its combustion produces toxicants that accumulate in smoke, filters and ashes^5^. More than 80% percent of these chemical compounds are inhaled into human lungs through cigarette smoking, some of which are known to be toxic or even carcinogenic^5,6^. These compounds are associated with more than 50 diseases that have the potential to cause damage to human cells and tissues, specifically increased risks of cancer, respiratory and cardiovascular diseases have been reported^7,8^. Earlier studies found that cigarette smoking is a risk factor related to increased prescriptions of antibiotics to smokers^9,10^. In the context of AMR, this increased consumption of antibiotics can contribute to the proliferation of ARGs in human associated microbial communities, as exposure to antibiotics, even at very low concentrations can result in the selection for ARGs^11–13^. Here we hypothesize that, aside from this indirect effect through increased antibiotic consumption, the toxic compounds accumulating in smoke, filters and ashes possess the potential to also directly affect the spread of AMR in the human lung as well as the environment.

One of the main drivers underlying the spread of ARGs is horizontal gene transfer (HGT) within and between bacterial populations, which is mediated by mobile genetic elements (MGEs) including plasmids, transposons, and phages^14–16^. Among these, the conjugative transfer of ARG-encoding plasmids is the most relevant in the context of AMR^17,18^. While the rate of plasmid transfer in typical natural settings is low, it can be significantly upregulated when bacteria are exposed to chemical stressors including antibiotics themselves^19,20^, but also non-antibiotic pharmaceuticals, heavy metals or nanomaterials^21–24^. The upregulation in genes connected with increased plasmid transfer and uptake is regularly observed for compounds triggering the bacterial stress response by causing an increase in intracellular reactive oxygen species (ROS)^25^. With cigarette smoke containing thousands of chemical compounds, many of which possess toxic properties^5,6^, we consider it likely that cigarette smoke can equally trigger stress responses in bacteria of the human lung microbiome. This could in turn result in elevated plasmid transfer rates and hence an increase in resistant bacteria, which could negatively affect antibiotic treatments of subsequent bacterial lung infections. To test this we here perform plasmid transfer experiments between fluorescently-tagged *Pseudomonas putida* KT2440 donor and recipient strains^16^ in artificial lung mucus medium, under exposure to different concentrations of cigarette smoke condensate. We further explore the underlying mechanisms of the observed increase in plasmid transfer rates under smoke exposure.

Aside from the lung, significant amounts of cigarette waste products enter the environment. Cigarette filters, added as a protective device, can trap chemical compounds and reduce the immediate health risk of smoking. However, with the huge consumption of cigarettes, around 1.2 million tons of filters per year are produced and discarded, together with the entrapped toxicants^26^, making them an important environmental pollutant. Cigarette filter leachates have for example been proven toxic to marine and freshwater fish^27^. The final waste product, cigarette ash, also contains a multitude of hazardous compounds such as the heavy metals arsenic, lead, cadmium and nickel, as well as organic toxicants, which can also leach into the environment^28,29^. Consequently, we propose that toxic leachates from cigarette filters and cigarette ash can, similar to smoke in the lung microbiome, increase plasmid transfer rates in environmental microbiomes. To test this, we performed plasmid transfer experiments, as described above, in the presence and absence of different concentrations of cigarette filters and cigarette ash leachates in liquid media.

Importantly, filters can persist in the environment for a long time due to their poor biodegradability^30^. With their high surface area^31^, filters could additionally provide perfect breeding grounds for high-risk microbes such as antibiotic resistant pathogens, especially since entrapped toxicants could provide co-selective pressure for increased colonization of bacteria hosting ARGs^32^, while simultaneously leading to increased HGT rates among these resistant colonizers. With their small size and light weight, colonized filters can easily be transported from wastewaters to various environments^33–35^ which might allow bacteria that colonize them to hitchhike to novel environments, a phenomenon previously described for microplastics^36^. Cigarette filters could hence not only increase the local abundance of ARGs, but also support their dissemination across habitat boundaries into novel microbiomes. To test if communities colonizing used cigarette filters are indeed enriched in high-risk microbes, we submerged used and unused cigarette filters in a wastewater stream in Dresden, Germany for five weeks and explored how the entrapped toxicants alter the colonizer microbiome with regard to the relative abundance of ARGs, mobile genetic elements and pathogens.

We hence provide novel insights into how cigarette consumption and the spread of antimicrobial resistance in both, human and environmental microbiomes, are immediately linked.

## Results

### Cigarette smoke promotes plasmid transfer by triggering the bacterial stress response

To test if cigarette smoke affects the plasmid-mediated transfer of antibiotic resistance genes (ARGs) in the smoker lung, mating experiments in artificial lung sputum medium were performed between the donor strain, *Pseudomonas putida* KT2440::*mCherry* hosting plasmid pKJK5::*gfpmut3b*, and the recipient strain, a rifampicin-resistant mutant of *Pseudomonas putida* KT2440. Mating experiments were executed in the absence of and under exposure to three different cigarette smoke condensate (CSC) concentrations. During the 24 hours mating experiments, successful growth of both, donor and recipient was observed in the sputum medium across all concentrations of CSC. The CSC concentrations did not significantly affect the final concentration of either the donor or the recipient strain (All *P* ≥ 0.064, ANOVA) (Fig. 1A & B). However, the final transconjugant concentration for every CSC exposure condition tested was significantly higher than in absence of CSC (*P* < 0.05) (Fig. 1C). The final density of transconjugants was positively correlated with the CSC concentration (r_s_ = 0.797, *P* < 0.001, Spearman) and reached up to 5.16 ± 2.19 × 10^7^ CFU mL^-1^ at the highest concentration of CSC (0.25 cigarette equivalents mL^-1^), accounting for a more than 3-fold increase in plasmid receipt (Fig. 1D & E). Similarly, the ratio of transconjugants normalized by the number of recipients, increased significantly with increasing CSC concentrations (r_s_ = 0.506, *P* = 0.012, Spearman), from 0.14 ± 0.11 to 0.38 ± 0.19 (Fig. 1E). To explore the underlying mechanisms of the CSC-induced promotion of plasmid transfer, the level of intracellular reactive oxygen species (ROS) production in *P. putida* was determined flowcytometrically. ROS production has regularly been reported as an indicator of bacterial stress, a main factor in increased plasmid transfer rates in bacterial populations^37^. Here, ROS production was significantly increased under exposure to CSC at all concentrations (All *P* < 0.05, ANOVA) by 1.24-1.46-fold compared to the control (Fig. 1F). This illustrates one among several potential pathways regarding how CSC exposure can increase the frequency of ARG transfer in the lung of smokers by triggering the bacterial stress response.

**Figure 1:**
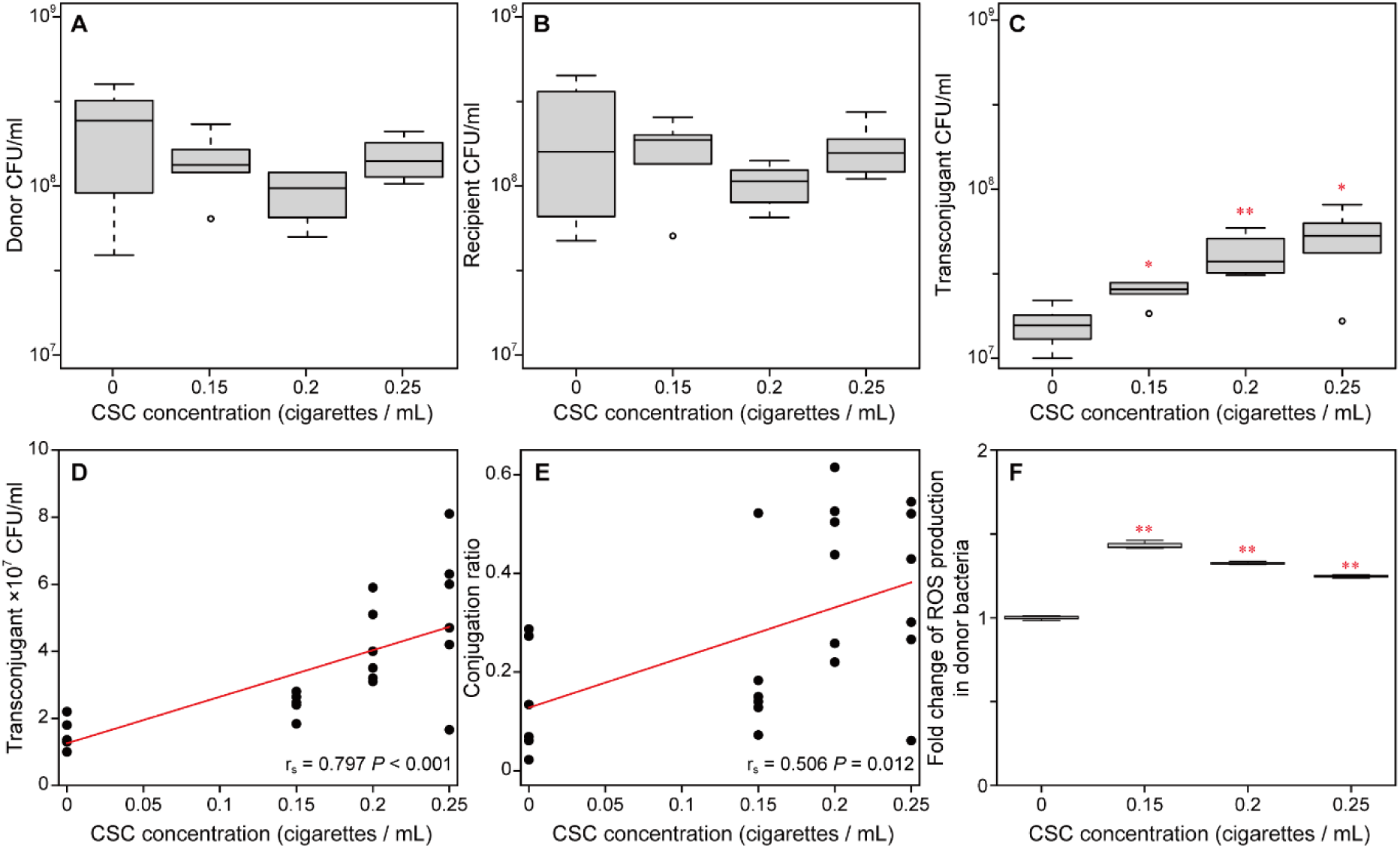
Effect of cigarette smoke condensate (CSC) on plasmid transfer. Final concentrations of **(A)** donor strain *P. putida* with plasmid pKJK5, **(B)** recipient stain *P. putida*, **(C)** transconjugant. Positive Spearman’s correlation of the cigarette smoke condensate concentration and **(D)** final concentration of the transconjugant and **(E)** conjugation ratio displayed as transconjugants per recipient. The linear fit line (red line) corresponding to Spearman’s correlation, spearman’s correlation coefficient r_s_ and statistics significance *P* are shown. **(F)** Fold change of ROS production in donor bacteria. Significant differences between treatments and control are indicated with stars based on ANOVA testing. *: *P* < 0.05; **: *P* < 0.01. The sample size for each treatment was n = 6.

### Effect of cigarette ash and cigarette filter leachate on plasmid transfer

In addition to direct effects in the lung, discarded cigarette ash and cigarette filters could equally promote ARG plasmid transfer in the environment. To test if this is the case, mating experiments were repeated in liquid growth medium in the presence and absence of different concentrations of cigarette ash solution (CAS) and cigarette filter leachate (CFL). Again, no significant reduction of bacterial growth was detected at any concentration of CAS or CFL (All *P* ≥ 0.475, ANOVA) (Fig. 2A & B, Fig. S1A & B). For CAS, similar to CSC, the density of transconjugants significantly increased by 4-fold from 8.19 ± 2.83 × 10^5^ CFU mL^-1^ in the absence to 3.49 ± 1.20 × 10^6^ CFU mL^-1^ at 0.025 cigarette equivalents mL^-1^ of CAS (*P* < 0.01) (Fig. 2C). A significant positive correlation between CAS concentration and the absolute number of transconjugants (r_s_ = 0.853, *P* < 0.001, Spearman, Fig. 2D) as well as the normalized ratio of transconjugants per recipients (r_s_ = 0.774, *P* < 0.001, Spearman, Fig. 2E) was observed. Contrary, no significant effects on plasmid transfer could be detected for the CFL, even if the concentration of cigarette equivalents was increased to two times that of the CAS (Fig. S1C). The intracellular production of ROS significantly increased under exposure to CAS by 3.91 ± 0.12 folds at 0.0125 cigarette equivalents mL^-1^, and 5.88 ± 0.36 folds at 0.025 cigarette equivalents mL^-1^ CAS (All *P* < 0.01, ANOVA) (Fig. 2F), indicating that bacterial stress plays an important role in promoting plasmid transfer. The main compounds causing this stress response under CAS exposure are likely high concentrations of heavy metals leaching out (Fig. S2), while only low heavy metal concentrations were detected for CSC. However, organic stressors such as nicotine, PAHs and other hydrocarbons are expected to also play a major role^29^. For CFL, only low concentrations of heavy metals were leaching into the solution, and the lack of an increase in plasmid transfer or bacterial ROS production indicates that also toxic organic compounds are rather stably bound to the filters.

**Figure 2:**
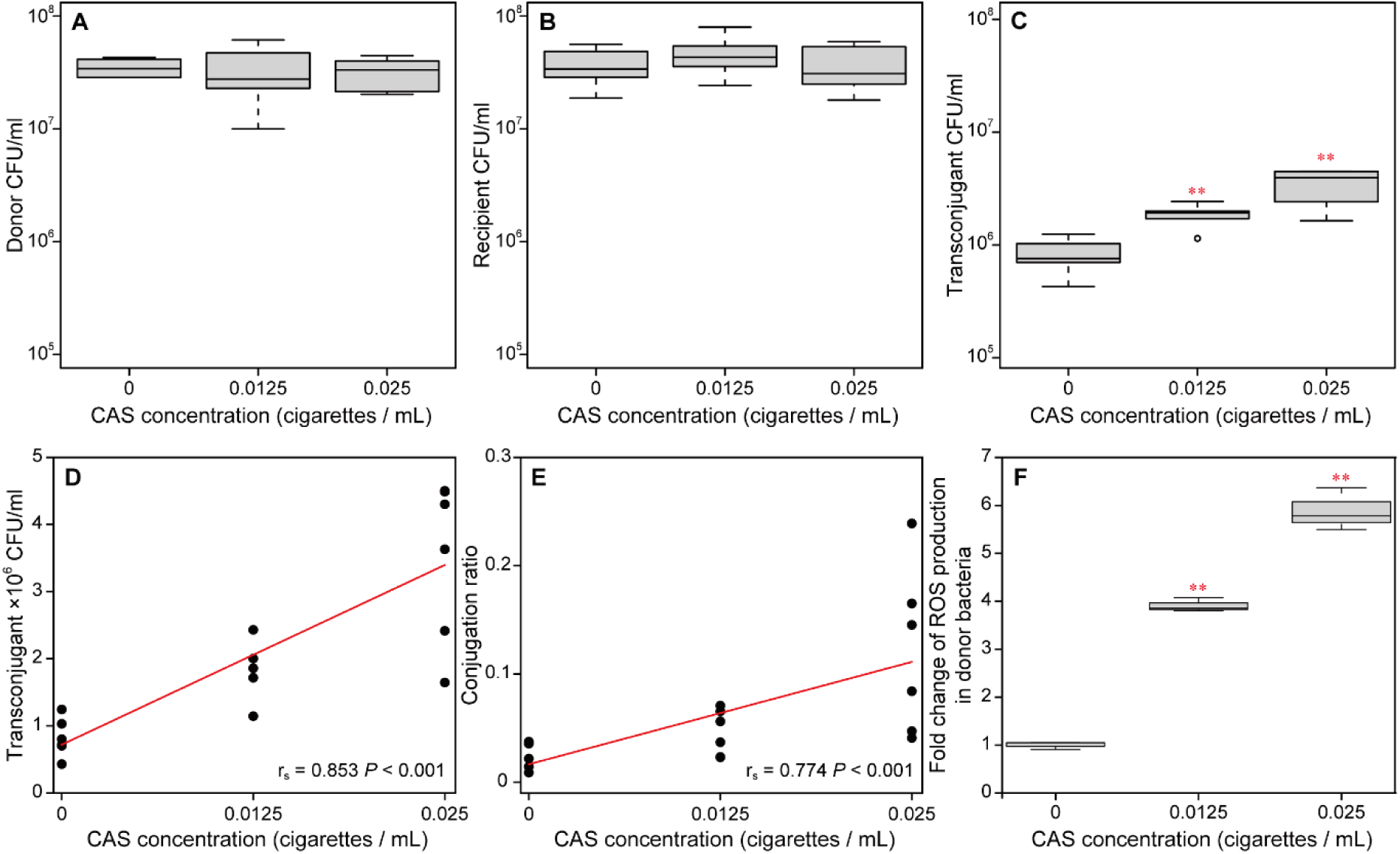
Effect of cigarette ash solution (CAS) on plasmid transfer. Final concentrations of **(A)** donor strain *P. putida* with plasmid pKJK5, **(B)** recipient stain *P. putida*, **(C)** transconjugant. Positive Spearman’s correlation of the cigarette ash solution concentration and **(D)** final concentration of the transconjugant and **(E)** conjugation ratio based on transconjugants per recipient. The linear fit line (red line) corresponding to Spearman’s correlation, spearman’s correlation coefficient r_s_ and statistics significance *P* are shown. **(F)** Fold change of ROS production in donor bacteria. Significant differences between treatments and control are indicated with stars based on ANOVA testing. *: *P* < 0.05; **: *P* < 0.01. The sample size for each treatment was n = 6.

### Used cigarette filters are colonized by a distinct microbial community

The toxic compounds bound to cigarette filters after cigarette smoking rarely leached into aqueous solution in the previous experiments and a hence likely to be rather stably bound to the filter. Still, high amounts of cigarette filters with these stably bound toxicants end up in wastewater^38^ and are regularly released from there to aquatic environments^27^. Hence, we questioned if these filters could provide breeding grounds bacterial communities with high levels of resistance adapted to the stably bound toxic compounds and in turn be vehicles on which high risk bacteria that colonize these filters could hitchhike to novel environments. To test this, we submerged 12 used filters, created using our automated smoking device, containing the toxic smoke compounds and 12 unused control filters in a wastewater stream for 5 weeks and thereafter analyzed the colonizing microbial communities. Both types of filters were successfully colonized by bacteria, with no significant difference in the number of colonizers being observed based on absolute 16S rRNA gene copies between used (9.44 ± 5.25 × 10^8^) and unused filters (1.77 ± 0.75 × 10^9^; *P* = 0.56, n = 24; t-test; Fig. S3). However, used filters were colonized by a significantly distinct microbial community compared to unused filters (AMOVA, *P* < 0.001, n = 24, Fig. 3A). In addition, the variation in the colonizing microbial community composition was far higher based on average Bray-Curtis dissimilarity between samples among the used (0.38 ± 0.09) compared to the unused filters in the NMDS plot (0.21 ± 0.02; *P* < 0.001, n = 24; ANOVA; Fig. 3A). On the phylum level, both groups of filters were mainly colonized by the same four phyla (Fig. 3B). However, the proportions significantly shifted on used filters towards increases in Proteobacteria (22.5 ± 6.3% on used vs. 16.7 ± 1.8% on unused, *P* < 0.01, ANOVA), Bacteroidetes (14.2 ± 5.2% vs. 8.8 ± 1.4%, *P* < 0.01) and Actinobacteria (6.1 ± 2.4% vs. 4.5 ± 0.8%, *P* = 0.05) at the cost of a decrease in the most dominant Firmicutes (34.3 ± 5.6% vs. 42.8 ± 1.6%, *P* < 0.001) (Fig. 3B). Among the rarer phyla, no significant differences between used and unused filters were observable (Fig. S4). Together this indicates that the embedded toxicants have indeed an effect on colonization and are potentially heterogeneously distributed among and on the filters causing the increased variation in colonizers.

**Figure 3:**
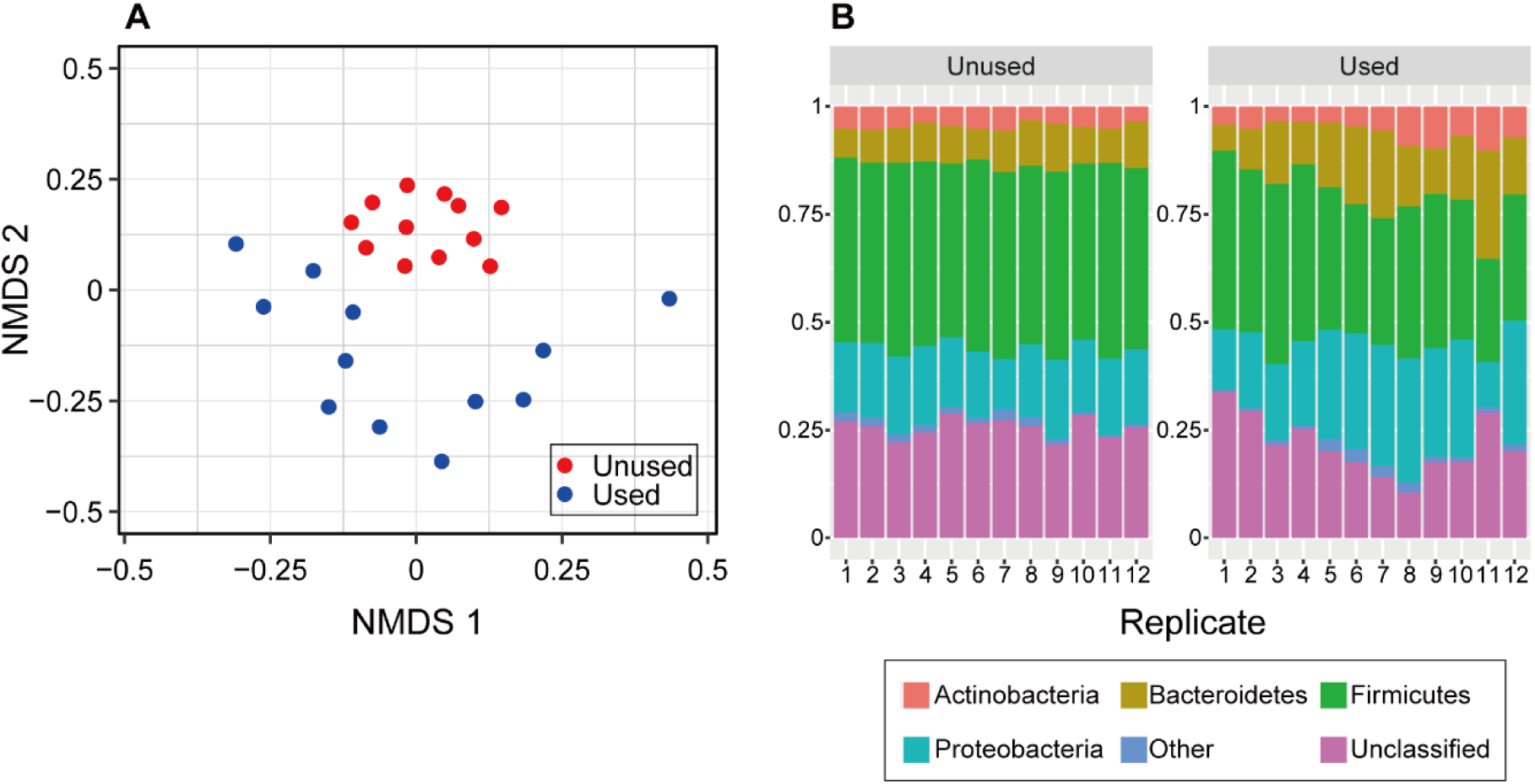
Microbial community composition of colonizers from a wastewater stream on used and unused filters. **(A)** NMDS beta-diversity plot based on Bray Curtis dissimilarity between samples at the 97% OTU level. **(B)** Dominant phyla composition on the 12 replicate unused and used filters. Phyla with an average relative abundance below 1% are shown as “Others” and are displayed in higher resolution in Figure S3.

To assess the potential pathogenicity of the colonizing bacteria on used and unused filters, their genus level distribution, identified through 16S sequence analysis, was compared to a comprehensive list of currently reported bacterial human pathogens^39^. Used filters were significantly more colonized by bacteria belonging to genera in which previously human pathogens have been reported (22.8 ± 3.9%) than unused filters (16.2 ± 1.5% *P* < 0.001, n = 24, ANOVA) (Fig. 4A). Consequently, a higher pathogenicity potential can be assumed. To confirm this trend, the relative abundance of the model pathogen *Klebsiella pneumoniae* was assessed using HT-qPCR. It was chosen as the model pathogen, as multi-drug resistant isolates of *K. pneumoniae* were previously reported as main pathogenic colonizers of plastic particles in the environment^40^. *K. pneumoniae* was found on used filters at significantly increased relative abundance of 4.7 ± 3.2 × 10^-5^ per 16S compared to the unused control filters (2.5 ± 1.7 × 10^-5^, *P* < 0.001, n = 24, ANOVA, Fig. 4B).

**Figure 4:**
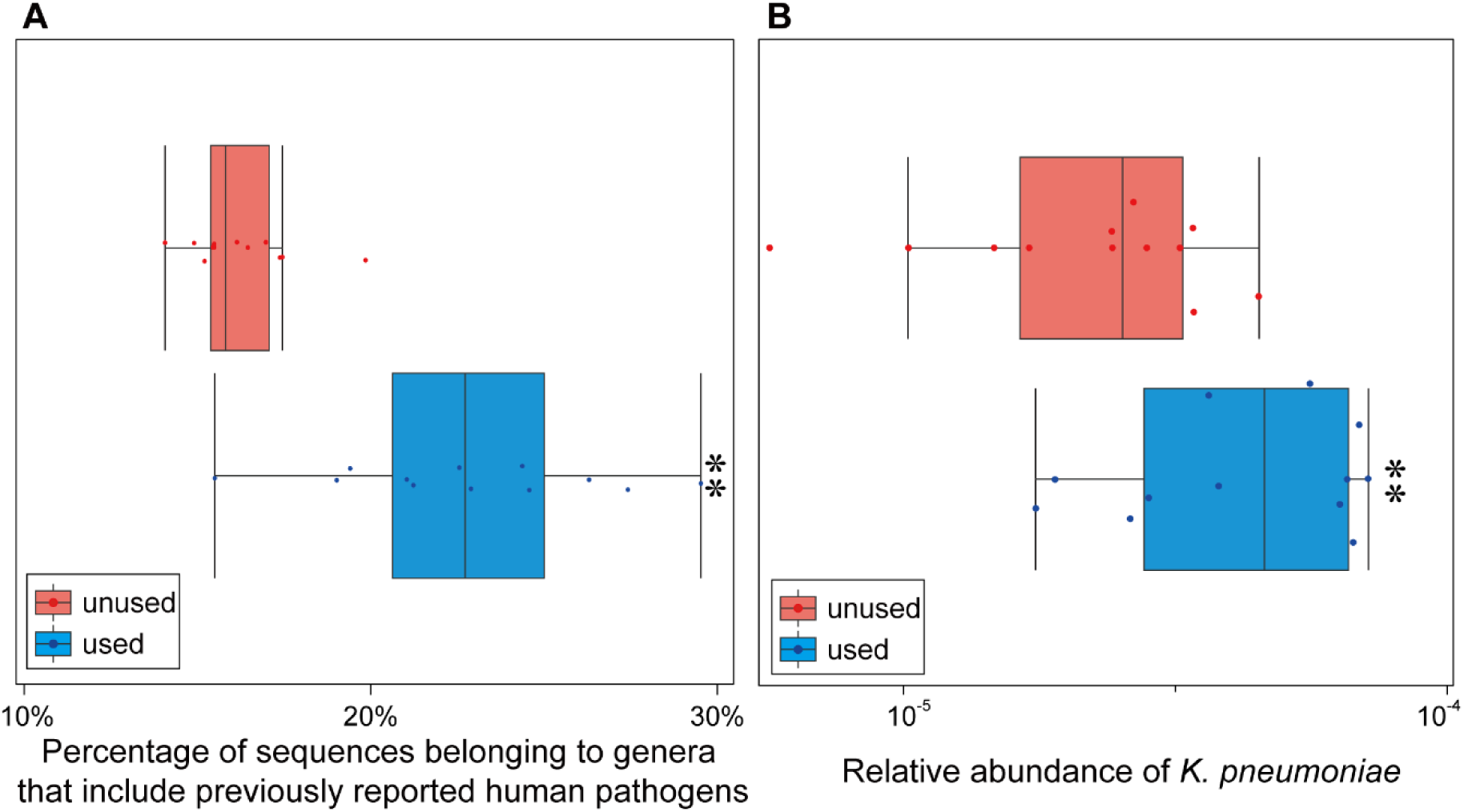
Pathogenic potential of the colonizing microbiome on used and unused cigarette filters. **(A)** Percentage of sequences belonging to genera that include previously reported human pathogens based on Bartlett et al.^39^. **(B)** Relative abundance per 16S rRNA gene of the model pathogen *Klebsiella pneumoniae* in the microbial community. Points in boxplots refer to individual measurements of individual samples within a treatment. Significant differences between used and unused control filters are indicated with stars based on ANOVA testing. *: *P* < 0.05. **: *P* < 0.01. Sample size for each treatment was n = 12.

### Cigarette filters as hot-spots and vehicles for the spread of antimicrobial resistant pathogens

Used cigarette filters were not only colonized by a distinct microbial community, but this community also contained a higher proportion of ARGs per bacterium. 24 out of 26 surveyed ARGs had a significantly increased relative abundance in communities colonizing used versus unused filters (*P* < 0.05; n = 12; Fig. 5A) with the increases ranging between 20.4-559.4% based on high-throughput chip-based qPCR. These ARGs belonged to 10 different classes of antibiotics (e.g., aminoglycosides, beta-lactams, quinolones, etc.) providing evidence that bacteria with increased levels of ARGs are more likely to also thrive on filters with toxicants embedded in them. ARGs were not only more abundant, but likely also more mobile, as all five of the assayed genetic markers for mobile genetic elements had a significantly increased relative abundance on used filters (all *P* < 0.01; n = 36; Fig. 5B). Based on the increased ARG abundance, genetic mobility and pathogenic potential, bacterial communities of increased risk to human health colonize used cigarette filters and can potentially hitchhike on these filters to novel environments.

**Figure 5:**
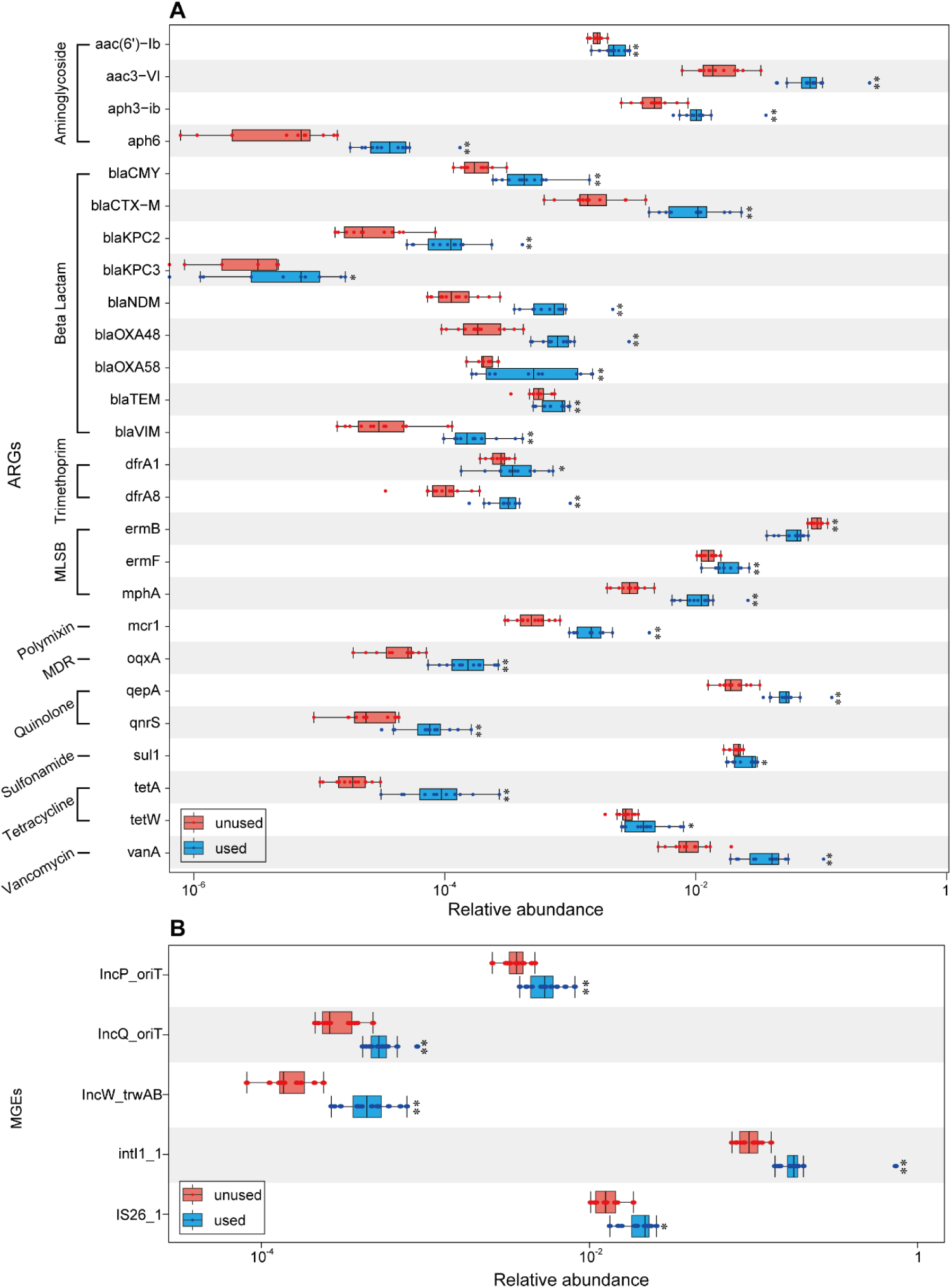
Relative abundance of **(A)** antimicrobial resistance genes (ARGs), **(B)** mobile genetic element marker genes per copy of the 16S rRNA gene in the colonizing microbiome on used and unused cigarette filters. Significant differences between used and unused control filters are indicated with stars based on ANOVA testing. *: *P* < 0.05; **: *P* < 0.01. The sample size for each treatment was n = 12.

## Discussion

In this study, we elucidate two independent mechanisms through which cigarette-derived compounds and waste products can facilitate the spread of AMR in the human lung as well as the environment. First, cigarette smoke and ash leachate were found to significantly promote horizontal transfer rates of a multidrug resistance plasmid. This coincided with an increased triggering of the bacterial stress response. While no effect of filter leachate on gene transfer was determined, used filters, enriched in toxicants and discarded in the environment, provided perfect breeding grounds for high-risk microbial communities that were significantly enriched in ARGs, MGEs and potential human pathogens, which possess the potential to hitchhike on these filters to novel habitats. This demonstrates that the consumption of cigarettes has not only direct, but also indirect adverse effects on human health by promoting the dissemination of AMR in bacterial communities.

Cigarette smoke condensate promoted plasmid transfer by more than two-fold in artificial lung sputum medium^41^, which suggests that similar effects are also observable in the human lung microbiome. Importantly, it was previously demonstrated, that effects of stressors on HGT in single strain *P. putida* experiments^22^ are regularly transferable to the complex community level^42^. Further, with the average adult human lung hosting about 30-100 mL of sputum that is renewed approximately every 24 hours^43^, the lowest observed effect concentration in our study of 0.15 cigarette equivalents of CSC per mL suggests that smoking 4.5-15 cigarettes per day would be sufficient to reach concentrations that trigger elevated plasmid transfer rates in the lung. However, this estimate is likely conservative as a) the exchange surface of up to 70 m² and exchange time of several seconds for toxicant transfer between smoke and lung mucus are far increased compared to the here used bubbling method for obtaining CSC^44^ and b) heterogeneity in the uptake of toxicants in lung mucus could lead to locally elevated concentrations that were avoided in our well-mixed experimental setup.

Experiments were here carried out using *Pseudomonas putida*, regularly recognized as an opportunistic pathogen causing lung infections^45–47^. Moreover, *Pseudomonas putida* has been reported to act as a reservoir of multidrug resistance elements in clinical settings from which transfer of ARGs to *Pseudomonas aeruginosa*^48^, an even more important lung pathogen, is regularly realized. Hence, the here described promotion effect of CSC on plasmid transfer in the lung of smokers could increase the prevalence of multi-antibiotic resistant opportunistic pathogens. This could result in higher difficulty of treatments and an elevated antibiotic treatment intensity^49^ should a subsequent infection occur.

One of the main underlying mechanism of cigarette smoke and ash leachate promoting plasmid transfer was the induction of an overproduction of reactive oxygen species (ROS), which can cause damage to cells, further altering membrane function and enhancing membrane permeability^50^. Oxidative stress caused by ROS can also activate the SOS stress response which regulates gene expression, including those involved in horizontal gene transfer^51,52^. Triggering of the stress response and ROS overproduction leading to increased plasmid transfer rates has previously been reported for heavy metals, non-antibiotic pharmaceuticals, and antibiotics^19,51,53^. Based on genotoxic effects of cigarette smoke condensate on *Pseudomonas* spp.^9,41^, and various toxic compounds detected in cigarette smoke, ashes, and filters including heavy metals and polycyclic aromatic hydrocarbons (PAHs)^21,29^ the here observed increased ROS production in response to cigarette-derived compounds is in line with expectations and can explain the realized promotion of plasmid transfer.

Similar effects regarding HGT were observed for cigarette ash leachate, however, with the low amount of ash discarded compared to other compounds selecting for and promoting HGT of ARGs^54^, the environmental effect size of this pollutant is likely to be small. Contrary, the around 1.2 million tons of low-degradable cigarette filters discarded per year provide a significant environmental pollutant^26^. As cigarette filters effectively capture a range of harmful and toxic substances, they become reservoirs of toxicants that can exert selective pressure on microbial communities^55^.

We here demonstrate that this selection pressure leads to the enrichment of bacterial taxa that possess metabolic capabilities or tolerance mechanisms to cope under such stressful conditions. The resulting communities originating from wastewater were, compared to unused control filters, enriched in ARGs, MGEs and pathogens. Pathogenic bacteria are known to possess various mechanisms that allow them to thrive in stressful conditions^56,57^ with the accumulation of toxicants in used filters potentially creating niches that selects for such bacteria with higher stress tolerance. Additionally, as discussed above, HGT of MGEs is increased under stress conditions in microbial communities^42,58^, leading to the observed enrichment in MGEs and their encoded ARGs on used filters that might serve as a hot-spot of genetic exchange. Additionally, the presence of toxicants can facilitate direct selection or co-selection of ARGs and MGEs, wherein ARGs either directly confer resistance to the present toxicants or are encoded on the same genetic element as genes that confer a fitness advantage^32^, leading to their proliferation within the filters. Further, PAHs and other aromatic hydrocarbons on cigarette filters could enrich degradation-relevant genes, which, if encoded on MGEs, could potentially co-select for ARGs as well^59^.

The potential dissemination of used cigarette filters enriched with potentially harmful bacteria from wastewater^60^ poses hence a significant concern for human and environmental health. Filters can persist in aquatic environments for up to 10 years^27^ and travel long distances. The microbial communities enriched in resistant pathogens can hence be dangerous hitchhikers to novel habitats, where they can potentially impact local ecosystems and spread infectious diseases to human or wildlife populations, a process previously described for microplastic particles^36^. This emphasizes the urgent need for proper cigarette filter disposal practices to mitigate this risk.

In conclusion, we here elucidated two different ways through which two of the biggest threats to human health are immediately connected. Cigarette smoking amplified the threat of AMR through promotion of plasmid transfer and creation of highly selective environmental niches for AMR and pathogens on cigarette filters.

## Material & Methods

### Cigarette smoke condensate (CSC)

Cigarette smoke condensate (CSC) was obtained following the protocols of Alamri^61^ and Abdelmalek et al.^41^, with minor modifications. In this study, the cigarette (Marlboro Red, Philip Morris International, Virginia, USA) was connected to a vacuum pump supplying a constant suction of 100 mbar. The cigarette was then lit and the resulting smoke was drawn through two sequential bottles containing 50 mL of sterile H_2_O each. The procedure was repeated for a total of 10 cigarettes. Then the two 50 mL solutions containing the smoke condensate compounds were combined to reach a 0.1 cigarette equivalent mL^-1^ CSC solution. The resulting CSC was further up-concentrated by evaporation using an aspirator pump to a final concentration of 1 cigarette equivalent mL^-1^ of water. The obtained CSC was sterilized by filtration through a 0.22 μm pore size sterilized filter (Millipore) and stored at 4 °C for use in subsequent experiments.

### Cigarette ash solution (CAS)

To prepare the cigarette ash solution (CAS), we collected the ashes of 10 cigarettes and added 100 mL sterile NaCl solution (0.9%). The mixture was incubated at 37 °C with shaking at 150 rpm for 24 h to allow ash compounds to leach into the solution. The final CAS at a concentration of 0.1 cigarette equivalents mL-1 was sterilized by filtration through a 0.22 μm pore size sterile filter (Millipore) and stored at 4 °C for use in subsequent experiments.

### Cigarette filter leachate (CFL)

Equally the used filters of 10 cigarettes were added to 100 mL sterile NaCl solution (0.9%). The cigarette filter solution was incubated at 37 °C with shaking at 150 rpm for 24 h to allow filter-bound compounds to leach into the solution. Thereafter the resulting cigarette filter leachate (CFL) at a final concentration of 0.1 cigarette equivalents mL^-1^ was sterilized through a 0.22 μm pore size sterile filter (Millipore) and stored at 4 °C for use in subsequent experiments.

### Metal analysis

The heavy metal content (cadmium, chromium, copper, manganese and zinc) of the resulting solutions was carried out with ICP-OES (optical emission spectrometry with inductively coupled plasma) Spectra Avio 200 (Perkin Elmer LAS GmbH, Rodgau, Germany) according to DIN standard EN ISO 11885. For this purpose, each sample was acidified with nitric acid (0.5 mL 65 % nitric acid in 100 mL sample) and disintegrated in a microwave (Mars CEM GmbH, Dresden, Germany). Disintegration was carried out with approximately 0.5 g raw sample with 5 mL distilled water and 5 mL concentrated nitric acid + 1 mL hydrogen peroxide. Subsequently, samples were filtered in 50 mL volumetric flasks through paper filters, topped up with deionized water and then measured with ICP-OES (DIN EN ISO 11885).

### Artificial sputum medium

To mimic the environment in the human lung, artificial sputum medium (ASM) was prepared according to the recipe in Palmer et al.^62^. Detailed information of the ingredients is provided in Table S1. Besides the components described in the previous study, 5 mL sterilized egg yolk and 5 g sterilized mucin were added per 1 L of sterile ASM to mimic the natural viscosity of lung mucus as previously suggested by Sriramulu et al.^63^.

### Bacterial strains

*Pseudomonas putida* KT2440::*mCherry* carrying the plasmid pKJK5::*gfpmut3b* was used as the donor strain for plasmid transfer experiments. *Pseudomonas putida* has been chosen as the model focal strain as it is known to cause lung infections, has been recognized as an opportunistic pathogen and is equally commonly found in diverse environments^45–47^. The donor strain was chromosomally tagged with a gene cassette *lacI_q_-pLpp*-mCherry*-Km^R^*, which encodes kanamycin (Km) resistance, *mCherry*-induced red fluorescence, and constitutive *lacI_q_* production that suppresses the expression of green fluorescent protein (GFP) on the plasmid. The plasmid pKJK5::*gfpmut3b* encodes tetracycline (Tet) and trimethoprim resistance^64^ and was tagged with the *gfpmut3* gene, which results in the expression of GFP in the absence of lacIq production^16^ upon transfer to the recipient. A rifampicin (Rif) resistant mutant of the wild-type *Pseudomonas putida* KT2440 strain^65^ was used as the recipient strain in this study. Expression of GFP is repressed in the donor stain but released in the recipient strain upon successful plasmid transfer, resulting in green fluorescence in transconjugant cells. As a result, the donor, recipient, and transconjugant can be selected for based on differing antibiotic resistances on solid media and differential fluorescence profiles under the fluorescence microscope.

### Plasmid transfer experiments

To mimic the environment in the human lung, plasmid transfer experiments between the donor and recipient strains in the presence of CSC were conducted in an artificial sputum medium (ASM). Contrary, plasmid transfer experiments in the presence of CAS and CFL were conducted in the Luria-Bertani medium. The donor and recipient strains were incubated overnight at 30°C with shaking at 150 rpm and corresponding antibiotics added (50 μg mL^-1^ Km and 10 μg mL^-1^ Tet for the donor, 100 μg mL^-1^ Rif for the recipient), harvested, washed and adjusted to OD_600_ = 1. Donor and recipient were then mixed at a 1:1 ratio, and added to 2 mL of the respective mating medium at a final concentration of around 10^6^ CFU mL^-1^. Thereafter CSC, CAS and CFL were added at the appropriate concentrations. CSC was added at final concentrations of 0, 0.15, 0.2, and 0.25 cigarette equivalents mL^-1^. CAS was added at 0, 0.0125, and 0.025 cigarette equivalents mL^-1^. CFL was added at 0, 0.025, and 0.05 cigarette mL^-1^. Six biological replicates were conducted for each concentration. After 24 h of incubation at 30 °C with shaking at 150 rpm, the mating mixtures were harvested, serial diluted in sterile 0.9% NaCl solution and plated on selective LB agar plates for enumeration of donor, recipient and transconjugant cells. The donor strain was selected for with Km 50 μg mL^-1^ and Tet 10 μg mL^-1^, the recipient strain was selected for with Rif 100 μg mL^-1^, and the transconjugants were selected for with Tet 10 μg mL^-1^ and Rif 50 μg mL^-1^. All plates were subsequently incubated at 30°C for 48 h, colonies counted, and checked under a fluorescence microscope for the correct expression of fluorescence. The conjugation rate was calculated by normalizing the number of transconjugants by the number of recipients.

### Measurement of intracellular reactive oxygen species (ROS) production

ROS production of the donor strain was measured by using a 2’,7’–dichlorofluorescein diacetate (DCFDA) kit (abcam®, UK), following the manufacturer’s protocol. The identical method was previously used for similar purposes^42^. Briefly, cell suspensions harvested in the exponential phase at a concentration of about 10^6^ CFU mL^-1^ were incubated with 20 μM DCFDA at 37 °C for 30 min in the dark. Then the cell suspensions were washed and incubated with the respective concentrations of CSC, CAS and CFL for 4 h in the dark. Both positive (150μM Tert Butyl Hydroperoxide) and negative control were included. All cell suspensions were then evaluated using a BD Accuri C6 flow cytometer (BD Accuri; Ann Arbor, MI, USA) with the following setting parameters (Threshold 500 on FL1-H, Fluidics Medium, Core Size 16 µm, Flow Rate 35 µl min^-1^, Run Limits 25 µl) and excitation at 488 nm to emission at 535 nm to obtain the average fluorescence per cell (defined by size through forward and side scatter) under the respective conditions. By dividing the average fluorescence obtained for each treatment by the fluorescence obtained for the negative control, the fold change in ROS production per cell can be measured as the DCFDA is converted into its fluorescent form through reaction with intracellular ROS. Increased ROS production consequently results in increased fluorescence.

### Colonization experiment with used and unused cigarette filters

Twelve replicate used and unused cigarette filters were fixed with fishing lines in individual 3D printed torpedo style housings that allow water to stream through while minimizing the risk of ragging due to wastewater compounds (e.g., toilet paper)^66^. These housings were then submerged into a wastewater stream at a monitoring site (51.011°N, 13.840°E) within a combined sewer network that is part of an Urban Observatory (Institute of Urban Water Management, TU Dresden) for 5 weeks. Following this, the filters were transported to the laboratory and washed twice with a sterile 0.9% NaCl solution to prevent residual wastewater bacteria from being attached to the filter. Then DNA was extracted from the filter colonizing community using the DNeasy PowerWater kit (Qiagen, Hilden, Germany) following the manufacturer’s protocol. The quality and quantity of extracted DNA were evaluated using a NanoDrop^TM^ (ThermoScientific, Waltham, MA, USA), and DNA was stored at -20 °C for downstream analysis.

### Quantitative PCR analysis to determine absolute bacterial abundance on colonized filters

To determine the absolute bacterial abundance on used and unused filters, qPCR for the 16S rRNA gene as an indicator for the total microbial abundance was performed using the following primer pair (Fw: TCCTACGGGAGGCAGCAGT, Rev: ATTACCGCGGCTGCTGG)^67^. The reactions were performed in technical triplicates on a MasterCycler RealPlex (Eppendorf, Germany) at a final volume of 20 μL with 10 μL of Luna Universal qPCR Master Mix (New England Biolabs, Germany), which uses SYBR Green chemistry. The template volume was 5 μL. The amount of DNA per reaction was standardized to 10 ng. The thermal cycling was operated with one initial denaturation cycle of 95 °C for 10 min followed by 40 cycles consisting each of 15 s at 95 °C and 1 min at 60 °C. Standard curves with amplification efficiency 0.9–1.1 and R^2^ ≥ 0.99 were accepted. Melting curve analysis was performed to assess the amplicons’ specificity. No PCR inhibition was detected based on serial dilution of the template DNA.

### Molecular analysis of the filter colonizing microbiome

To analyze the microbial diversity and taxonomic composition of the samples, DNA extracts were sent to the IKMB sequencing facility (minimum 10,000 reads per sample; Kiel University, Germany). Illumina MiSeq amplicon sequencing of the bacterial 16S rRNA gene was performed using primers targeting the V3-V4 region (V3F: 5′-CCTACGGGAGGCAGCAG-3′ V4R: 5′-GGACTACHVGGGTWTCTAAT-3)^68^. Sequencing analysis was executed using the Mothur software package v.1.48.0^69^ following the MiSeq SOP^70^. Taxonomy assignment was then performed on resulting OTUs (97%) to classify the sequences into taxonomic groups, with a focus on phylum-level composition. All sequencing data were submitted to the NCBI sequencing read archive under project accession number PRJNA1002237.

### Analysis of potential pathogenicity

The 16S rRNA sequences were processed on the latest version of the QIIME2 (Quantitative Insights Into Microbial Ecology) platform (https://qiime2.org/)^71^. First of all, raw sequencing data was imported into QIIME2 and demultiplexed, and primers were cut with the command ‘cutadapt’. Quality control and denoising of the data were further performed using DADA2^72^, and the feature table was obtained at ASV level. Finally, there was an average of 10, 935 reads per sample in the feature table. The representative sequences were annotated using a pre-trained Naïve Bayes classifier based on the SILVA database (v138) on QIIME2. The data was further compared to a comprehensive list to calculate the percentage of sequences belonging to genera that include human pathogens^39^.

### High-throughput qPCR analysis of resistome, mobile genetic elements and pathogen abundance

Additionally, to study the relative abundance of ARGs, mobile genetic elements and pathogens, the extracted DNA was sent to Resistomap Oy (Helsinki, Finland) for high-throughput qPCR analysis using a Smart chip real-time PCR system. In total, 33 primer sets^73^ were utilized targeting the abundance of 26 antibiotic resistance genes, 5 mobile genetic element (MGE) marker genes, 1 taxonomic marker for the pathogen *Klebsiella pneumoniae* and 16S rRNA gene (Table S2). The protocol was as follows: PCR reaction mixture (100 nL) was prepared using SmartChip TB Green Gene Expression Master Mix (Takara Bio, Shiga, Japan), nuclease-free PCR-grade water, 300 nM of each primer, and 2 ng/μL DNA template. After initial denaturation at 95 °C for 10 min, PCR comprised 40 cycles of 95 °C for 30 s and 60 °C for 30 s, followed by melting curve analysis for each primer set. A cycle threshold (Ct) of 31 was selected as the detection limit^74,75^. Amplicons with non-specific melting curves or multiple peaks were excluded. The relative abundances of the detected gene to 16S rRNA gene were estimated using the ΔCt method based on mean CTs of three technical replicates^76^.

### Statistics

Spearman’s correlation coefficients and significance of correlation were calculated in SPSS v22.0 (IBM Corp, Armonk, NY, USA). One-way analysis of variance (ANOVA) was performed in SPSS v22.0, and significant differences between groups were calculated by the Tukey post hoc test. Distance between samples based on microbial community composition was calculated based on the Bray-Curtis dissimilarity metric^77^. Significant grouping of samples in the NMDS plot were assessed based on analysis of molecular variance (AMOVA)^78^.

## Acknowledgments

UK, DK, RPM, GO, PK & TUB were supported by the Urban Resistome project funded by the Deutsche Forschungsgemeinschaft (DFG) under project number 460816351. PF was supported through the China Scholarship Council (CSC) under grant number 202004910327. UK, TUB, JH & BL were supported by the ExploreAMR project funded by the Bundesministerium für Bildung und Forschung under grant number 01DO2200. AE & EF were supported through the ACRAS-R and the PRESAGE project funded by the Bundesministerium für Bildung und Forschung under grant numbers 16GW0355 and 02WAP1619. The authors thank C. Zschornack, S. Kunze, J. Isler and H. Brückner for technical support in the laboratories. Responsibility for the information and views expressed in the manuscript lies entirely with the authors.

## Data Availability

The datasets supporting the conclusions of this article are included within the article and its additional files or available through the corresponding author upon reasonable request. Original sequencing data is available in the NCBI sequencing read archive under project accession number PRJNA1002237.

## Author Contributions

PF, DKo, TUB & UK conceived the study. PF, AT-O & AD developed and optimized the artificial lung mucus medium for experiments. PF & EF performed plasmid transfer experiments. PF & GO performed the ROS assays. RPM & PK performed the heavy metal analysis. DKo & RPM performed the filter colonization experiments. DKo, AE, UK, JH & BL performed the bioinformatic analysis of the microbial communities and pathogen potential. PF, DKo, DKn & UK analyzed, interpreted and visualized the data. PF, DKo & UK wrote the initial draft of the manuscript. All authors edited the manuscript. The final manuscript has been approved by all authors.

## Supplementary information

This article is supplemented with three additional figures and two additional tables in the supplementary information.

