## Supplementary Information for "Cigarette smoking promotes the spread of antimicrobial resistance in the human lung and the environment"

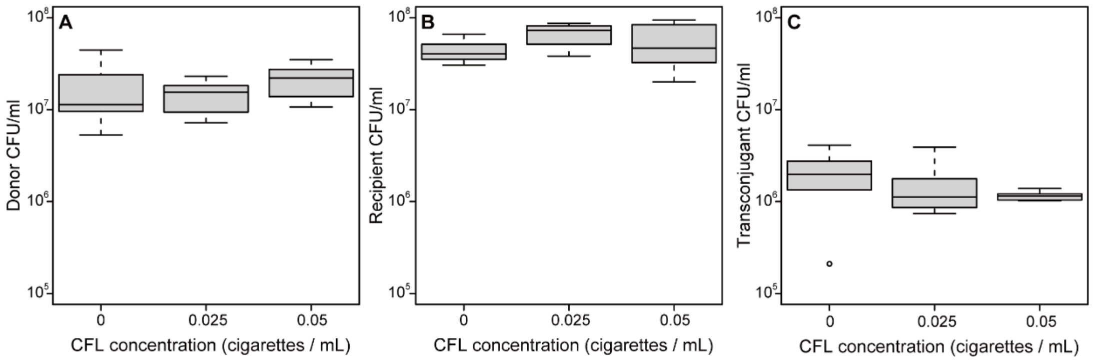


**SI Fig. 1:** Effect of cigarette filter leachate (CFL) on plasmid transfer. Final concentrations of **(A)** donor strain *P. putida* with plasmid pKJK5, **(B)** recipient stain *P. putida*, and **(C)** transconjugants. Significant differences between CFL concentration and control are indicated with stars based on ANOVA testing. *: $P < 0.05$; **: $P < 0.01$. Sample size for each treatment n = 6.

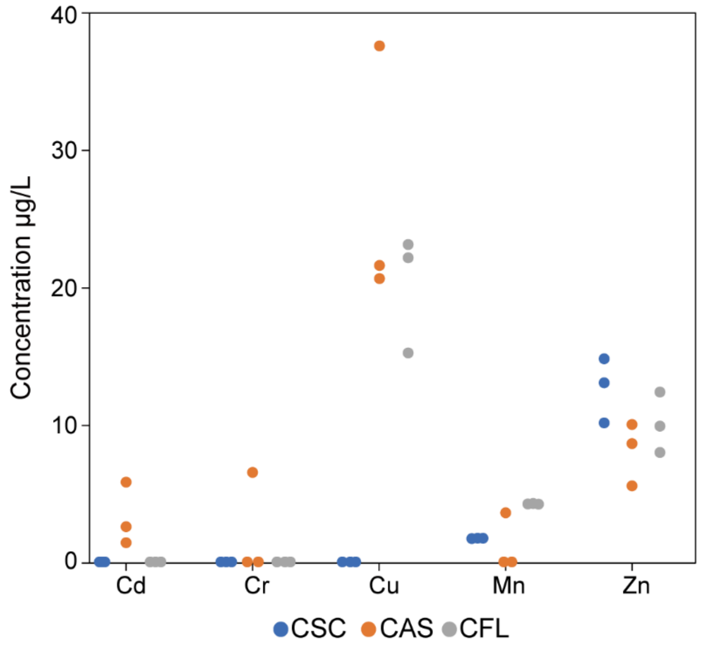


**SI Fig. 2:** Heavy metal concentrations in cigarette smoke solution (CSC), cigarette ash leachate (CAS) and cigarette filter leachate (CFL). Average concentrations from three individually prepared replicate samples are shown.

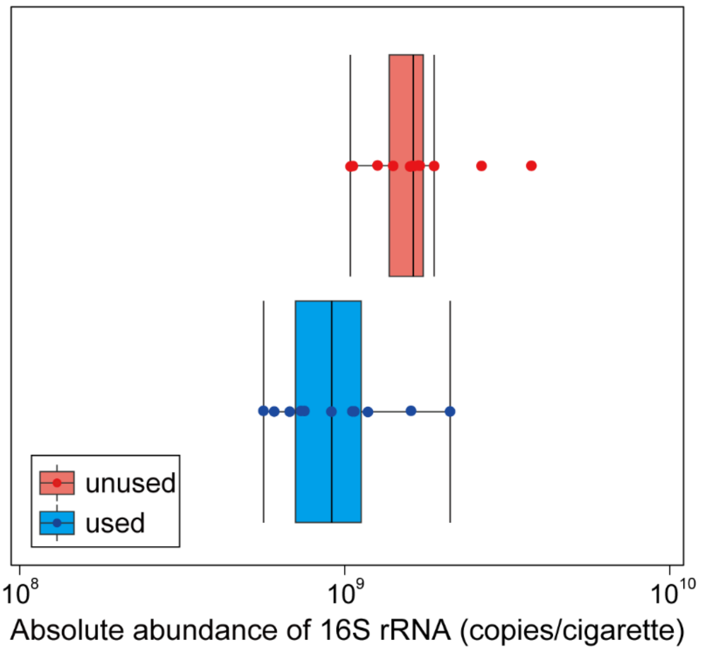


**SI Fig. 3:** Absolute bacterial abundance on used and unused cigarette filters after colonization in wastewater based on the 16S rRNA gene. Sample size for each treatment was n = 12. No significant difference was observed.

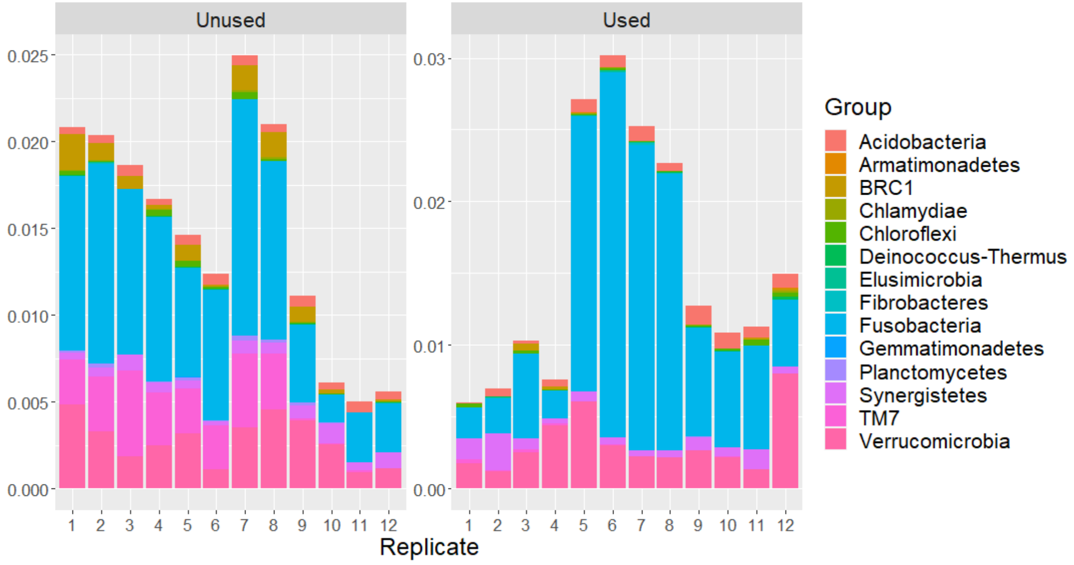


**SI Fig. 4:** Microbial community composition of colonizers from a wastewater stream used and unused filters based on phyla. In depth look into rare phyla with an average relative abundance below 1% that were grouped as “Others” in Figure 3.

**Table S1.** Components of artificial sputum medium.

| Components | Concentration (M) | Volume for 1L (mL) |
|---|---|---|
| l-aspartic acid sodium salt (in NaOH 0.5M) | 0.1 | 8.27 |
| l-threonine | 0.1 | 10.72 |
| l-serine | 0.1 | 14.46 |
| L-glutamic acid (in HCl 1M) | 0.1 | 15.49 |
| l-proline | 0.1 | 16.61 |
| l-glycine | 0.1 | 12.03 |
| l-alanine | 0.1 | 17.8 |
| l-cysteine | 0.1 | 1.6 |
| l-valine | 0.1 | 11.17 |
| l-methionine | 0.1 | 6.33 |
| l-isoleucine | 0.1 | 11.2 |
| l-leucine | 0.1 | 16.09 |
| l-tyrosine (in NaOH 1M) | 0.1 | 8.02 |
| l-phenylalanine | 0.1 | 5.3 |
| l-ornithine•HCl | 0.1 | 6.76 |
| l-lysine•HCl | 0.1 | 21.28 |
| l-histidine•HCl | 0.1 | 5.19 |

| | | |
|---|---|---|
| l-tryptophan (in NaOH 0.2M) | 0.1 | 0.13 |
| l-arginine•HCl | 0.1 | 3.06 |
| $NaH_2PO_4$ | 0.2 | 6.5 |
| $Na_2HPO_4$ | 0.2 | 6.25 |
| $KNO_3$ | 1 | 0.348 |
| $K_2SO_4$ | 0.25 | 1.084 |
| MOPS | 0.01 | 1 |
| $CaCl_2$ | 1 | 1.754 |
| $MgCl_2$ | 1 | 0.606 |
| $FeSO_4*7H_2O$ | 0.0036 | 1 |
| glucose | 1 | 3 |
| lactate | 1 | 9.3 |

**Table S2.** qPCR primers used in this study based on Stedtfeld et al.[1].

| Gene/Marker | Target | Forward Primer | Reverse Primer |
|---|---|---|---|
| 16S rRNA | 16S rRNA | GGGTTGCGCTCGTTGC | ATGGYTGTCGTCAGCTCGTG |
| *aac(6')-Ib* | Aminoglycoside | GTTTGAGAGGCAAGGTACCGTAA | GAATGCCTGGCGTGTTTGA |
| *aac3-VI* | Aminoglycoside | CGTCACTTATTCGATGCCCTTAC | GTCGGGCGCGGCATA |
| *aph3-ib* | Aminoglycoside | AACAGGTTTGGGAGGCGATG | CGCAACAAGCCTCTCCTGAA |
| *aph6* | Aminoglycoside | CCCATCCCATGTGTAAGGAAA | GCCACCGCTTCTGCTGTAC |
| *blaCMY* | Beta Lactam | AAAGCCTCAT GGGTGCATAAA | ATAGCTTTTGTTTGCCAGCATCA |
| *blaCTX-M* | Beta Lactam | CGTACCGAGCCGACGTTAA | CAACCCAGGAAGCAGGCA |
| *blaKPC2* | Beta Lactam | GCCGCCGTGCAATACAGT | GCCGCCCAACTCCTTCA |
| *blaKPC3* | Beta Lactam | CAGCTCATTCAAGGGCTTTC | GGCGGCGTTATCACTGTATT |
| *blaNDM* | Beta Lactam | GGCCACACCAGTGACAATATCA | CAGGCAGCCACCAAAAGC |

| | | | |
|---|---|---|---|
| *blaOXA48* | Beta Lactam | TGTTTTTGGTGGCATCGAT | GTAAMRATGCTTGGTTCGC |
| *blaOXA58* | Beta Lactam | GCAATTGCCTTTTAAACCTGA | CTGCCTTTTCAACAAAACCC |
| *blaTEM* | Beta Lactam | CGCCGCATACACTATTCTCAG | GCTTCATTCAGCTCCGGTTC |
| *blaVIM* | Beta Lactam | GCACTTCTCGCGGAGATTG | CGACGGTGATGCGTACGTT |
| *oqxA* | MDR | GAGTCAACCTACCTCCACTATCA | GCTGCGAGTTATCCAGCAG |
| *ermB* | MLSB | TAAAGGGCATTTAACGACGAAACT | TTTATACCTCTGTTTGTTAGGGAATTGAA |
| *ermF* | MLSB | CAGCTTTGGTTGAACATTTACGAA | AAATTCCTAAAATCACAACCGACAA |
| *mphA* | MLSB | CTGACGCGCTCCGTGTT | GGTGGTGCATGGCGATCT |
| *qepA* | Quinolone | GGGCATCGCGCTGTTC | GCGCATCGGTGAAGCC |
| *qnrS* | Quinolone | CCACTTTGATGTCGCAGATCTTC | CCCTCTCCATATTGGCATAGGAAA |
| *sul1* | Sulfonamide | GCCGATGAGATCAGACGTATTG | CGCATAGCGCTGGGTTTC |
| *tetA* | Tetracycline | GCTGTTTGTTCTGCCGGAAA | GGTTAAGTTCCTTGAACGCAAACT |

| | | | |
|---|---|---|---|
| *tetW* | Tetracycline | ATGAACATTCCCACCGTTATCTTT | ATATCGGCGGAGAGCTTATCC |
| *dfrA1* | Trimethoprim | GGAATGGCCCTGATATTCCA | AGTCTTGCGTCCAACCAACAG |
| *dfrA8* | Trimethoprim | GGTCGCACCTGCATCGTTA | AGCGCCACCAATGACGTAG |
| *vanA* | Vancomycin | GGGCTGTGAGGTCGGTTG | TTCAGTACAATGCGGCCGTTA |
| *mcr1* | Other | CACATCGACGGCGTATTCTG | CAACGAGCATACCGACATCG |
| *intI1* | MGE | CGAACGAGTGGCGGAGGGTG | TACCCGAGAGCTTGGCACCCA |
| IncP_oriT | MGE | CAGCCTCGCAGAGCAGGAT | CAGCCGGGCAGGATAGGTGAAGT |
| IncQ_oriT | MGE | TTCGCGCTCGTTGTTCTTCGAGC | GCCGTTAGGCCAGTTTCTCG |
| IncW_trwAB | MGE | AGCGTATGAAGCCCGTGAAGGG | AAAGATAAGCGGCAGGACAATAACG |
| IS26_1 | MGE | ATGGATGAAACCTACGTGAAGGTC | CGGTACTTAATCTGTCGGTGTTCA |
| *K. pneumoniae* | Taxonomic marker for pathogen | ACGGCCGAATATGACGAATTC | AGAGTGATCTGCTCATGAA |
